# Unannotated small RNA clusters in circulating extracellular vesicles detect early stage liver cancer

**DOI:** 10.1101/2020.04.29.066183

**Authors:** Johann von Felden, Teresa Garcia-Lezana, Navneet Dogra, Edgar Kozlova, Mehmet Eren Ahsen, Amanda J. Craig, Stacey Gifford, Benjamin Wunsch, Joshua T. Smith, Sungcheol Kim, Jennifer E. L. Diaz, Xintong Chen, Ismail Labgaa, Philipp K. Haber, Reena Olsen, Dan Han, Paula Restrepo, Delia D’Avola, Gabriela Hernandez-Meza, Kimaada Allette, Robert Sebra, Behnam Saberi, Parissa Tabrizian, Amon Asgharpour, Douglas Dieterich, Josep M Llovet, Carlos Cordon-Cardo, Ash Tewari, Myron Schwartz, Gustavo Stolovitzky, Bojan Losic, Augusto Villanueva

**Affiliations:** Division of Liver Diseases, Liver Cancer Program, Tisch Cancer Institute, Department of Medicine, Icahn School of Medicine at Mount Sinai, New York, NY, USA; Department of Internal Medicine, University Medical Center Hamburg Eppendorf, Hamburg, Germany; Department of Genetics and Genomic Sciences, Cancer Immunology Program, Tisch Cancer Institute, Icahn School of Medicine at Mount Sinai, New York, NY, USA; IBM T. J. Watson Research Center, Yorktown Heights, New York, NY, USA; Department of Visceral Surgery, Lausanne University Hospital CHUV, Lausanne, Switzerland; Department of Pathology, Icahn School of Medicine at Mount Sinai, New York, NY, USA; Liver Unit and Centro de Investigación Biomédica en Red de Enfermedades Hepáticas y Digestivas (CIBEREHD), Clínica Universidad de Navarra, Pamplona, Spain; Sema4, a Mount Sinai venture, Stamford, CT, USA; Department of Surgery, Icahn School of Medicine at Mount Sinai, New York, NY, USA; Liver Cancer Translational Research Laboratory, BCLC Group, IDIBAPS, CIBEREHD, Hospital Clinic, Universitat de Barcelona, Catalonia, Spain; Institució Catalana de Recerca i Estudis Avançats, Barcelona, Catalonia, Spain; Department of Urology, Icahn School of Medicine at Mount Sinai, New York, NY, USA; Icahn Institute for Data Science and Genomic Technology, Icahn School of Medicine at Mount Sinai, New York, NY, USA; Diabetes, Obesity and Metabolism Institute, Icahn School of Medicine at Mount Sinai, New York, NY, USA; Division of Hematology and Medical Oncology, Department of Medicine, Icahn School of Medicine at Mount Sinai, New York, NY, USA

**Keywords:** next-generation sequencing, genomics, cancer surveillance, liver cancer

## Abstract

**Background:** Hepatocellular carcinoma (HCC) is among the deadliest malignancies and surveillance tools for early detection are suboptimal. Extracellular vesicles (EVs) have gained increasing scientific interest due to their involvement in tumor initiation and metastasis, however, most extracellular RNA (exRNA) biomarker studies are limited to annotated genomic regions.

**Methods:** EVs were isolated with ultracentrifugation and nanoDLD and quality assessed by electron microscopy, immunoblotting, nanoparticle tracking, and deconvolution analysis. We performed genome-wide small exRNA sequencing, including unannotated transcripts. We identified small RNA clusters (smRCs) and delineated their key genomic features across biospecimens (blood, urine, tissue) and EV isolation techniques. A 3-smRC signature for early HCC detection was trained and validated in two independent cohorts.

**Results:** EV-derived smRCs were dominated by uncharacterized, unannotated small RNA and uniformly tiled across the genome with a consensus sequence of 20bp. A 3-smRC signature was significantly overexpressed in circulating EVs of HCC patients compared to controls at risk or patients with non-HCC malignancies (p<0.01, n=157). An independent validation in a phase 2 biomarker study revealed 86% sensitivity and 91% specificity for the detection of early HCC from controls at risk (i.e. cirrhosis or chronic liver disease, n=209) (positive predictive value (PPV): 89%, area under the ROC curve [AUC]: 0.87). The 3-smRC signature was independent of alpha-fetoprotein (p<0.0001) and a composite model yielded an increased AUC of 0.93 (sensitivity: 85%, specificity: 94%, PPV: 95%).

**Conclusion:** An exRNA-based 3-smRC signature from plasma detects early stage HCC, which directly leads to the prospect of a minimally-invasive, blood-only, operator-independent surveillance biomarker.

**One sentence summary:** We employ a novel, data-driven approach to identify and characterize small RNA clusters from unannotated loci in extracellular vesicle-derived RNA across different cancer types, isolation techniques, and biofluids, facilitating discovery of a robust biomarker for detection of early stage liver cancer.

## INTRODUCTION

Extracellular vesicles (EVs), including microvesicles and exosomes, are nanoparticles whose nucleic acid payload is capable of priming receptor cells to modify key cellular functions^1,2^. EVs are heterogeneous, both in terms of biogenesis and content^3^. While larger EVs such as apoptotic bodies mostly contain fragmented DNA, smaller EVs such as exosomes are enriched in non-coding, regulatory small RNAs (small RNAs)^2,4^. In cancer, EVs are increasingly recognized as key players in tumor initiation and metastasis^5^, mainly through miRNA trafficking, prompting their evaluation as early detection and treatment response biomarkers^6^. Most studies characterizing extracellular RNA (exRNA) and studying EV-related biomarkers apply conventional, reference-based, RNA sequencing approaches, and are thus limited to known annotated genomic regions. However, small RNAs arise from thousands of endogenous genes and are part of the genomic ‘dark matter’ of highly abundant yet largely uncharacterized non-coding RNA, with emerging roles in regulating gene expression via post-transcriptional and translational mechanisms. In fact, relatively little attention has been paid to characterizing the general expression landscape of circulating EV small RNA and their precursors in this context regardless of biotype, especially for those expressed from unannotated genomic regions.

Here, we adopt a different approach by *de novo* assembly and characterization of the small RNA expression landscape of exRNA, specifically including unannotated genomic regions. We define discrete loci called small RNA Clusters (smRCs), delineate their key properties, and test their clinical utility as early detection biomarkers in patients with liver cancer. Projections estimate more than 1 million deaths due to this cancer in 2030 worldwide^7^. With a 5-year survival of 18%, it is the second most lethal malignancy after pancreatic cancer. Survival in patients enrolled in early detection programs of hepatocellular carcinoma (HCC), the most common form of primary liver cancer, doubles that of those not enrolled in surveillance^8^. However, implementation of surveillance among patients at high risk of HCC in the United States is very low (20%)^9^ and the performance of recommended surveillance tools (i.e. ultrasound and serum alpha-fetoprotein (AFP)^10^) is suboptimal, with close to 40% of tumors being missed^11^. Using whole RNA sequencing of exRNA derived from EVs, we describe novel clinically-relevant smRCs in circulating EVs that can help detect HCC at an early tumor stage, which would allow patients to receive curative therapies.

## RESULTS

Our study is based on four independent cancer exRNA datasets: a prostate cancer cohort, which we termed the ‘smRC characterization’ cohort, to define and study the properties of smRCs in exRNA (9 patients, 41 samples), a ‘biomarker discovery’ cohort (n=15 patients) to identify differentially expressed smRCs between liver cancer patients and controls, an external dataset of patients with non-HCC malignancies (n=142) to test the HCC-specificity of our biomarkers, and an independent ‘biomarker validation’ cohort (n=209 patients) to confirm their clinical utility in a phase 2 biomarker study for detection of early stage HCC.

### Characterization of EV isolates

In all cohorts we employed differential ultracentrifugation (UC) to isolate exRNA, and followed the recommendations of the International Society of Extracellular Vesicles^12^ for quality assessment of EV isolates. Specifically, we used transmission electron microscopy, nanoparticle tracking analysis, immuno-labeling with Western Blotting for intracellular (i.e., tumor susceptibility gene 101 protein, TSG101) and Exoview™ for transmembrane (i.e., tetraspanins CD9, CD63, CD81) vesicle proteins in a subset of samples (**Fig. 1**). This suggested an enrichment for small EVs (median size of 120 nm) with compatible morphology and expression of typical markers. Additionally, for the ‘smRC characterization’ cohort in prostate cancer, we isolated exRNA from a subset of patients (n=5) using the ‘lab-on-chip’ technology nanoDLD^13^ (DLD) for serum samples. We also isolated purely cellular small RNA (<300 nt) from prostate cancer and adjacent non-cancerous tissue of the same patients to quantify exRNA-isolation technology, biofluid, and exRNA-specific variance in small RNA profiles, respectively. Part of our prostate cancer dataset has been included in an exRNA-atlas based deconvolution analysis published earlier^4^. An independent analysis found that our UC and nanoDLD isolation methods specifically isolate low- (cargo type 1) and variable (cargo type 4) density vesicles with minimum contamination from lipoproteins and argonaute proteins^4^. For this study, we have further performed the same computational deconvolution analysis for our ‘biomarker discovery’ dataset to determine carrier types and found that cargo type 4 was preferentially enriched. In fact, cargo type 4 is associated with vesicles in the 60 - 150 nm size range, which were purified consistently with nanoDLD, and also the lowest-density OptiPrep fractions 1-3 from serum and plasma^4^. Cargo type enrichments associated with low density vesicles, lipoproteins, AGO2-positive ribonucleoproteins (RNPs), and AGO-2 negative RNPs were significantly depleted (**Fig. 2**).

**Figure 1.**
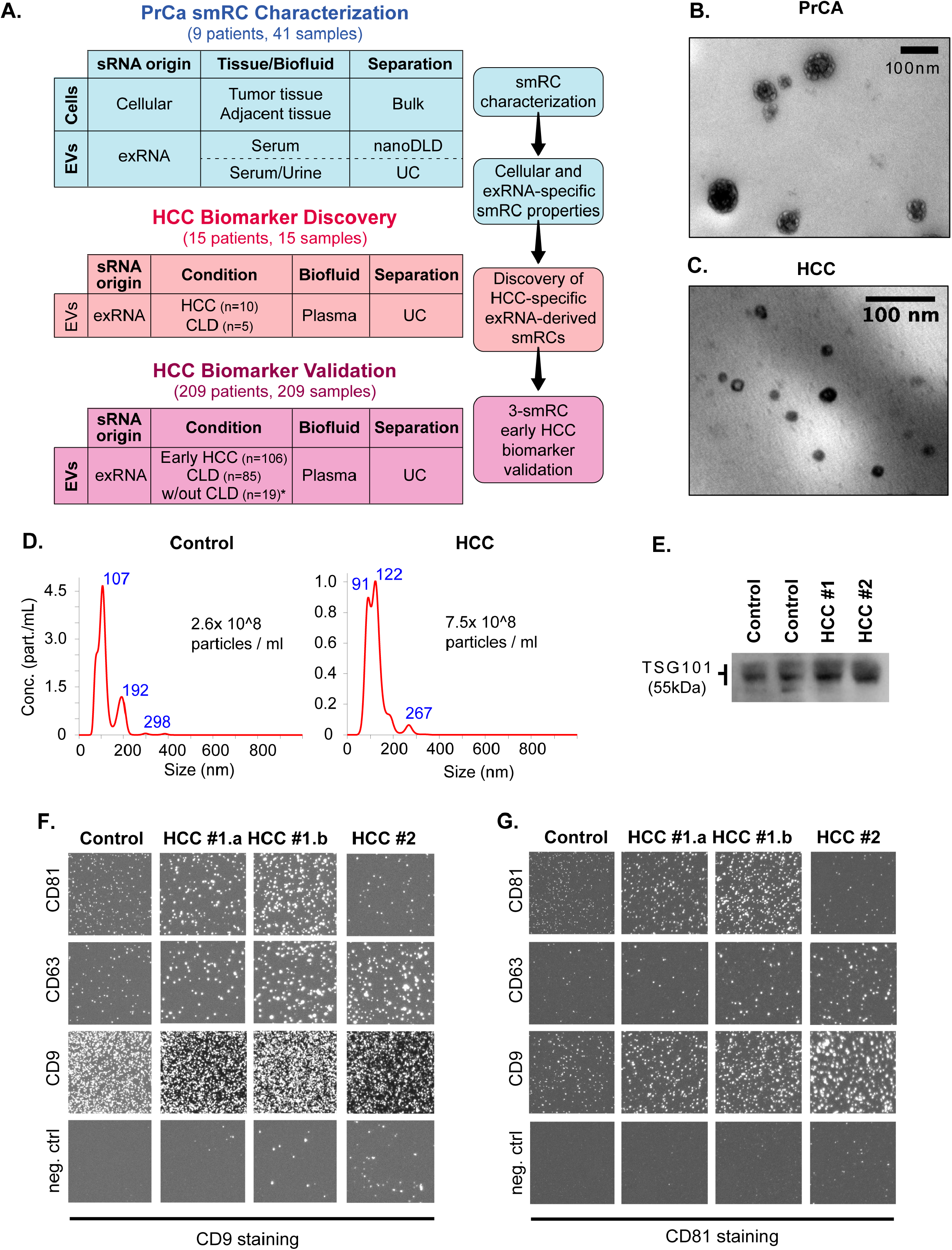
Summary and quality assessment of EV separation process for exRNA extractions from human plasma samples. (**A**) Schematic view of study flow diagram with different cohorts, and available specimen and separation analysis for each cohort. (**B, C**) Transmission electron microscopy image of prostate cancer serum isolate (B) and HCC plasma isolate (C). (**D**) Nanoparticle tracking analysis (Nanosight®) results in the plasma isolate of a control (left) and HCC (right) patient with corresponding size distribution and estimated particle concentration. (**E**) Western Blotting image of protein lysate from isolate against TSG101 (~55 kDa) in two control (left) and two HCC (right) patients. (**F, G**) Immunolabeling of the isolate with Exoview™. Isolates were captured by indicated antibodies (CD81, CD63, CD9, control IgG) on a chip and stained with CD9 (E) or CD81 (F) antibodies to visualize different EV subpopulations in one control and three HCC samples (#1.a and #1.b represent technical replicates from the same patient).

**Figure 2.**
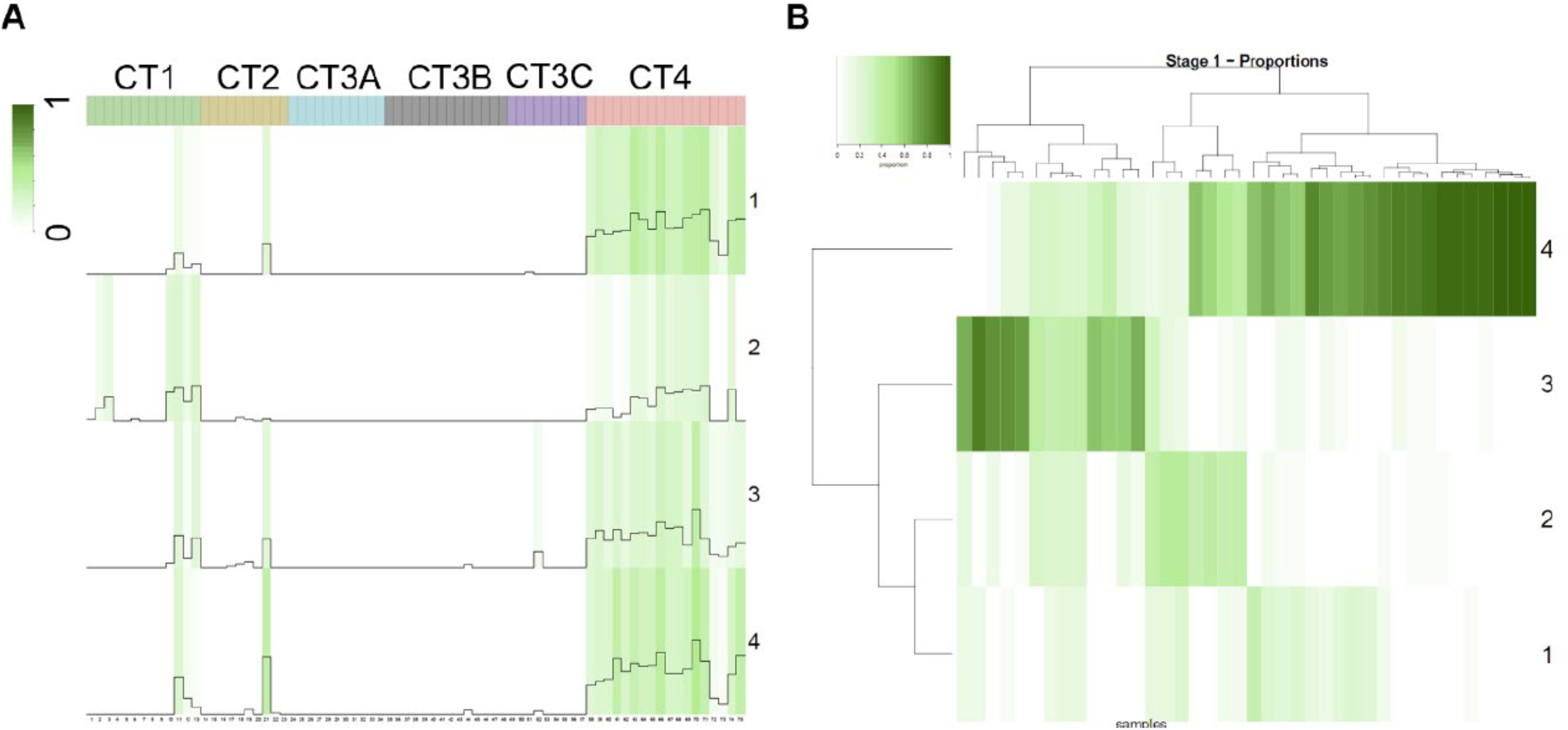
Annotated exRNA expression in cargo profiles. (**A**) Estimated constituent cargo profiles (rows) are correlated using the exRNA expression in transformed transcript abundance values across the informative RNAs against the 6 cargo types (CTs, columns) previously identified^4^. CT4 is heavily enriched, i.e. ncRNA profiles 58-75 are heavily enriched. (**B**) Per-sample proportions of estimated constituent cargo profiles. Heatmap of the per-sample proportions (columns) for each estimated constituent cargo profile (rows) numbered 1 through k.

### Identification and characterization of small RNA clusters from unannotated exRNA

We used small RNA sequencing data from the ‘smRC characterization’ and ‘biomarker discovery’ cohorts to define clusters of contiguous genomic regions with sufficient alignment coverage (termed ‘small RNA clusters’, smRCs). This allowed us to capture the known heterogeneous genome-wide expression of clusters of small RNA precursors^14^, each of which can give rise to multiple functional small RNA products, by defining clusters of small RNA reads (see **Fig. 3A**). Adjacent smRCs are merged if they overlap within a minimal padding threshold, and we define the key properties of smRCs: a) entropy (i.e., read tiling efficiency or complexity), b) peak coverage, and c) consensus sequence of each smRC. The set of all smRCs, computed once for all samples, is essentially the paired set of all accumulation loci of small RNA expression and their peak-coverage consensus sequences (which range from 15 to 100 nucleotides in length), and constitutes a smoothed, de-novo assembled small RNA expression landscape with a standard count matrix.

**Figure 3.**
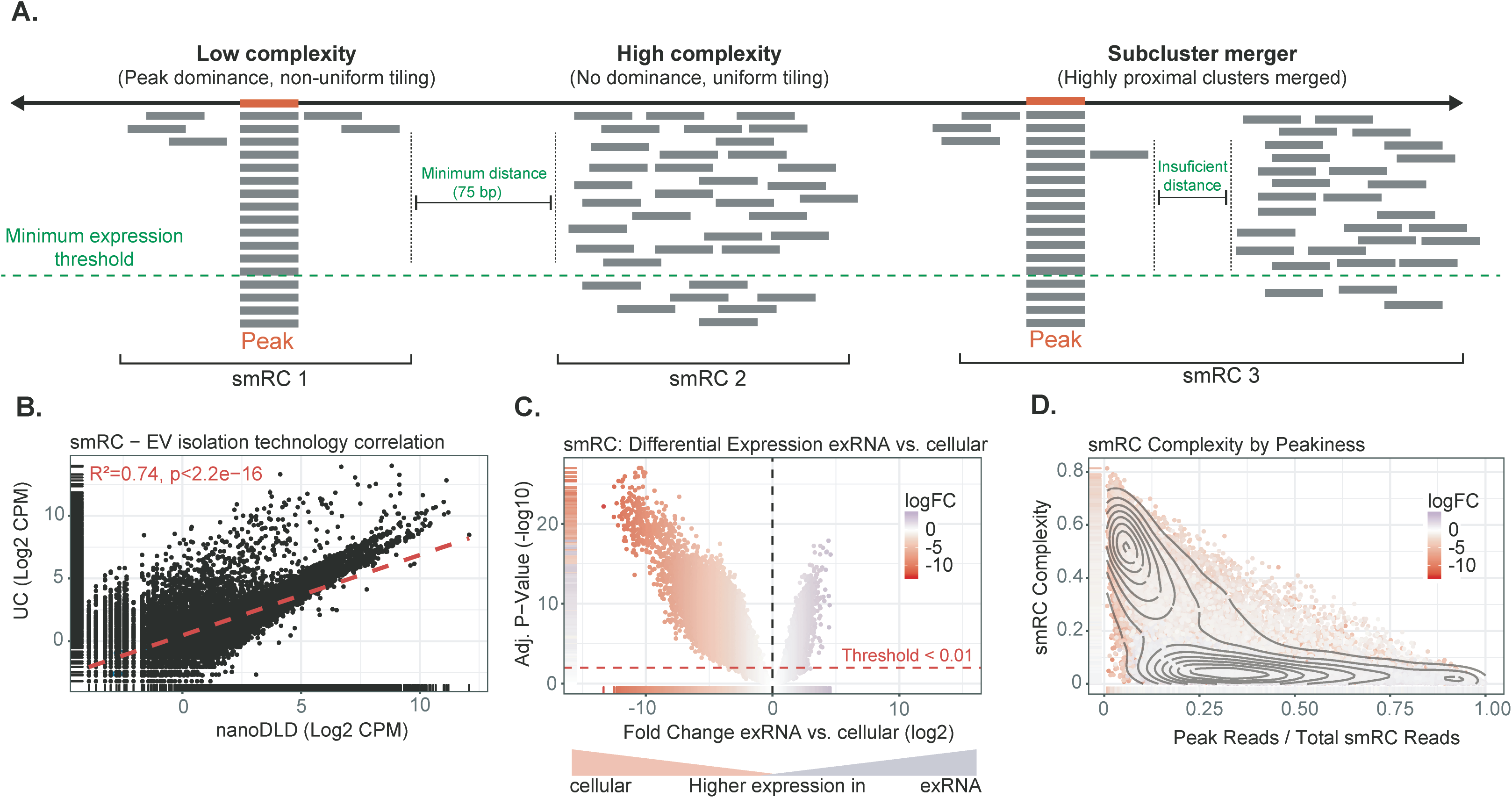
Key properties of small RNA clusters (smRCs). (**A**) Minimum coverage and sub-read length minimal spacing define smRCs. Read tiling complexity captures heterogeneity of smRC read distribution. (**B**) Correlation of smRC expression across different EV extraction methods (i.e., ultracentrifugation, UC, and nanoDLD). (**C**) Volcano plot for differential expression between smRC of cellular versus exRNA origin. (**D**) smRC complexity as a function of peak coverage colored by differential smRC expression between cellular and exRNA origin. smRCs enriched in exRNAs (purple) present with low complexity and higher peak coverage, whereas cellular smRCs (red) are more frequently of high complexity and lower peak coverage.

We delineated key genomic properties of smRCs in our ‘smRC characterization’ prostate cancer dataset due to the availability of different biological sample types (blood, urine, tumoral and non-tumoral adjacent tissue) and different isolation methods (ultracentrifugation and nanoDLD^15,16^). The mean genomic length of smRCs was 674 bp (**Supplementary Fig. S1A**), while the mean length of the consensus peak sequence was 20 bp. In order to profile the maximal coverage and overall distribution of expression within smRCs associated with exRNA, we defined two quantities. First, a ‘peak’ coverage which is simply the ratio of reads in the smRC peak to total smRC coverage, and second, a tiling complexity measure which is the ratio of unique read nucleotide sequences to total smRC coverage. smRCs with high complexity are those with uniform tiling and few peaks (see **Fig. 3A**). We found that the major contributor of smRC variable expression was RNA origin (with low complexity typical in exRNA- *versus* high complexity typical of cellular smRC origin, **Supplementary Fig. S1C**). Technical reproducibility of smRC quantification included comparing two different EV enrichment methods in serum (UC and nanoDLD), and different biofluid compartments (urine and serum) of the same patients. We found a high correlation between enrichment methods (spearman R^2^ ~ 0.74, p < 2.2e-16, **Fig. 3B**) with over 80% of smRCs detected by both methods above the 20th percentile of expression (**Supplementary Fig. S2A**). We found a modest correlation between different biofluid compartments (i.e. urine and plasma) using UC (spearman R^2^ ~ 0.45, p < 1e-16, see **Supplementary Fig. S2B+C** for self-reproducibility). Well-expressed smRCs possessed a heteroscedastic count variance profile which facilitated usual differential expression analysis via linear modeling (**Supplementary Fig. S1B**). The total number and magnitude of overexpressed smRCs in cells was significantly higher than in exRNA (**Fig. 3C**). However, we observed a significant difference in the complexity of smRCs found in exRNA compared to cells (**Fig. 3D)**. Indeed, the bimodal pattern reveals a clear separation between cellular smRCs, which overwhelmingly have relatively high tiling complexity, and exRNA smRCs that have much stronger evidence for high relative peak coverages. The mean size of the peak within smRCs was slightly higher than the minimal trimmed read length, and was significantly different between exRNA-derived and cell-derived (16.5 bp *versus* 22.6 bp, p < 1 e- 16). exRNA-associated smRCs preferentially overlap unannotated small RNA species compared to cellular smRCs (**Supplementary Fig. S3, Supplementary Table S1**). Finally, we orthogonally validated the expression of three unannotated smRCs using RT-qPCR (**Supplementary Fig. S2D**). These data demonstrate that low-complexity, exRNA-associated smRCs preferentially capture non-coding small RNA compared to protein-coding RNA, but are also significantly enriched in unannotated genomic regions.

### Small RNA clusters from plasma detect curable primary liver cancer

Given their high biological and technological independent reproducibility, tractable statistical properties, and unique ability to discriminate concentrations of exRNA-specific small RNA, we computed the smRC profile of our ‘HCC biomarker discovery’ cohort of 15 patients, including 10 patients with HCC and 5 controls at risk for HCC matched for age, sex, and etiology of the underlying liver disease (**Supplementary Table S2**). We found that exRNA-derived smRCs were differentially expressed between HCC and controls. In fact, 250 smRCs were enough to distinguish them (**Supplementary Fig. S4**). This led us to hypothesize that smRCs could be useful tools for early HCC detection. We selected the three top differentially expressed and low-complexity smRCs (see Supplementary Methods) and orthogonally validated their differential expression in this ‘HCC biomarker discovery’ cohort using RT-qPCR. Pearson’s correlation coefficient was higher than 0.6 for all three smRCs when comparing data from small RNA sequencing and RT-qPCR (p<0.05, **Supplementary Fig. S4B-D**). The three smRCs were located in regions of chromosomes 3q, 8q (both unannotated intergenic region), and 10q (intronic region of *SGPL1*) (**Supplementary Table S4**).

To determine the clinical relevance of smRCs in exRNA, we designed a phase 2 biomarker study following the recommendations from the Early Detection Research Network (EDRN) from the National Cancer Institute^17^. In detail, we aimed at assessing the role of our 3-smRC signature as a novel early detection biomarker in HCC. Recommended surveillance tools for early HCC detection (abdominal ultrasound and AFP)^10^ have low sensitivity (63%) and moderate specificity (83%)^11^. Improvement in this area is urgently needed by developing better read-outs of oncogenesis and facilitating implementation of surveillance through minimally-invasive, operator-independent tools. Unlike many studies in this setting^18^, we only enrolled patients with HCC at an early stage (Barcelona Clinic Liver Cancer classification^7^ (BCLC) stage 0 or A), who can be cured with either surgery or ablation^7^. Crucially, our control cohort is the target population for HCC surveillance as defined in clinical practice guidelines^10,19^. We included 209 patients (n=105 treatment-naive, early stage HCC, n=85 control patients with chronic liver disease (CLD) enrolled in HCC surveillance, and n=19 individuals without chronic liver disease (non-CLD) (**Table 1**). Following EV enrichment from plasma and exRNA extraction, we confirmed significant overexpression of our 3-smRC signature in HCC patients compared to CLD controls with qRT-PCR (p<3e-5, **Fig. 4A,** see **Fig. 5D** for comparison with non-CLD patients). Additional analysis in a cohort of 142 patients with other malignancies further confirmed their HCC specificity (**Fig. 5E**). To leverage the collective power of all three smRCs to predict early HCC risk we built a logistic regression model to discriminate between early HCC patients and CLD controls (excluding patients without chronic liver disease). This allowed us to test if there is a well calibrated and predictive association between smRC expression and early HCC detection (**Fig. 4D**). We used penalized maximum likelihood techniques, bootstrap and cross-validation to estimate and control for model optimism^20^, RT-qPCR batch plate effects, and over-fitting of our 3-smRC early detection signature. The logistic regression model was well calibrated with a low mean absolute probability error (0.04) to predict early HCC (**Fig. 4B**), low Brier score (B = 0.15), high AUC (0.87), and high Gini mean difference in predicted log-odds between HCC and CLD patients (2.44) adjusted under bootstrap (n = 1,000) resampling (**Fig. 4D**). Predicted HCC risk via smRC expression can be visualized via a patient nomogram to provide an individual estimate of HCC risk (**Fig. 5A**). In order to estimate sensitivity and specificity measures at plausible decision points, we applied the logistic regression model to a 85/15 split of the biomarker validation set for training and testing respectively. Averaging over 1,000 iterations, we recovered 86% sensitivity and 91% specificity with a positive predictive value of 89% on average by maximizing the balanced accuracy of the test ROC curves (**Supplementary Fig. S5 and Fig. 4C+D**). The area under the ROC curve (AUC) for our 3-smRC model was 0.87. Finally, a likelihood ratio test between an AFP-only early HCC detection model and one incorporating both AFP and our 3-smRC early detection signature showed that our smRCs add significant predictive power to AFP alone (p<0.0001). As expected, AFP levels and expression of our 3-smRC signatures were not correlated (**Fig. 5B**), which suggest that both capture complementary signals for early HCC detection. Indeed, a blood-based composite model of our 3-smRC signature and AFP yielded an increased AUC of 0.93, lower Brier score of 0.11, and better test performance (85% sensitivity, 94% specificity, positive predictive value of 95%, **Fig. 3C+D**). We also confirmed the EV origin of our smRC signal, as the expression of smRC-48615 was significantly higher using EV-enriched isolates as opposed to EV-depleted plasma (n=30 patients, **Fig. 5C**).

**Table 1.** Clinical characteristics of early stage HCC patients and controls.

|  | **Early Stage HCC**<br>**(n=105)** | **CLD, risk for HCC**<br>**(n=85)** | **Healthy liver (n=19)** |
| --- | --- | --- | --- |
| **Age (years)** | 65.5 | 62 | 52 |
| **Sex (male)** | 82 (80%) | 47 (55%) | 6 (32%) |
| **Cirrhosis (Yes)** | 68 (67%) | 61 (72%) | 0 |
| **Bilirubin (mg/dL)** | 0.7 | 1.0 | 0.6 |
| **Albumin (g/dL)** | 3.7 | 3.9 | 3.7 |
| **Platelets (count/mm<sup>3</sup>)** | 152.5 | 122.5 | 243 |
| **Etiology** |  |  |  |
| HCV | 44 (43%) | 22 (41%) | n.a. |
| HBV | 18 (18%) | 15 (28%) | n.a. |
| Alcohol | 9 (9%) | 12 (22%) | n.a. |
| NASH | 13 (13%) | 2 (4%) | n.a. |
| Other | 18 (18%) | 3 (6%) | n.a. |
| **Tumor stage (BCLC)** |  |  |  |
| Very Early (Stage 0) | 22 (21%) | n.a. | n.a. |
| Early (Stage A) | 83 (79%) | n.a. | n.a. |
| **Single nodule** | 92 (90%) | n.a. | n.a. |
| **Largest nodule (cm)** | 2.9 | n.a. | n.a. |
| **AFP (ng/mL\*)** | 8.3 | 3.8 | ** |

Continuous variables are displayed as median. \*Upper limit of normal 9 ng/mL. \*\*available data not representative of group (n=3). AFP, alpha fetoprotein, BCLC, Barcelona Clinic for Liver Cancer, HBV/HCV, chronic hepatitis B/C, NASH, non-alcoholic steatohepatitis, n.a., not applicable.

**Figure 4.**
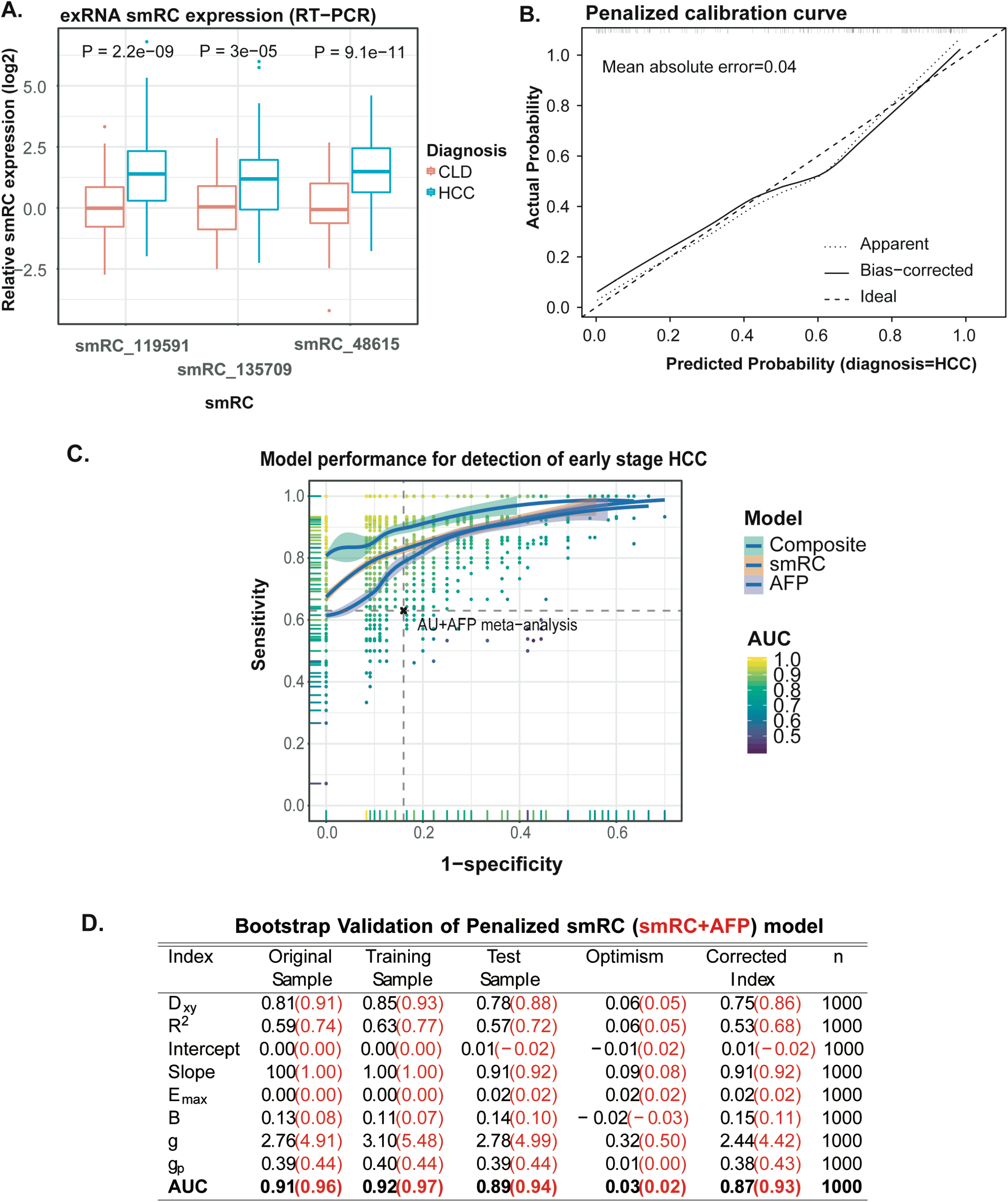
Performance of 3-smRC signature in a phase 2 biomarker study. (**A**) Expression for each smRC between HCC patients and chronic liver disease controls (CLD) (center line, median; box limits, upper and lower quartiles; whiskers, 1.5x interquartile range; points, outliers). (**B**) Calibration curve for penalized smRC logistic regression model to predict early HCC, with mean error (**C**) ROC curve for maximized gain-of-certainty across repeated cross validation. Each point represents a pair of sensitivities and specificities that maximize gain-in-certainty (i.e. sensitivity + specificity) from a test validation ROC curve, whose AUC colors the point. The loess curves trace the best density fit of points across this space, with 95% confidence intervals shown in gray. (**D**) Bootstrap validation parameters for smRC and smRC+AFP model. Dxy: Somers’ rank correlation between the observed HCC status and predicted HCC probabilities; Emax: maximum absolute calibration error on probability scale; B: Brier score; g: Gini’s mean difference of log-odds between HCC and CLD; gp: Gini’s mean difference in probability scale; AUC: Area Under the Receiver Operating Curve.

**Figure 5.**
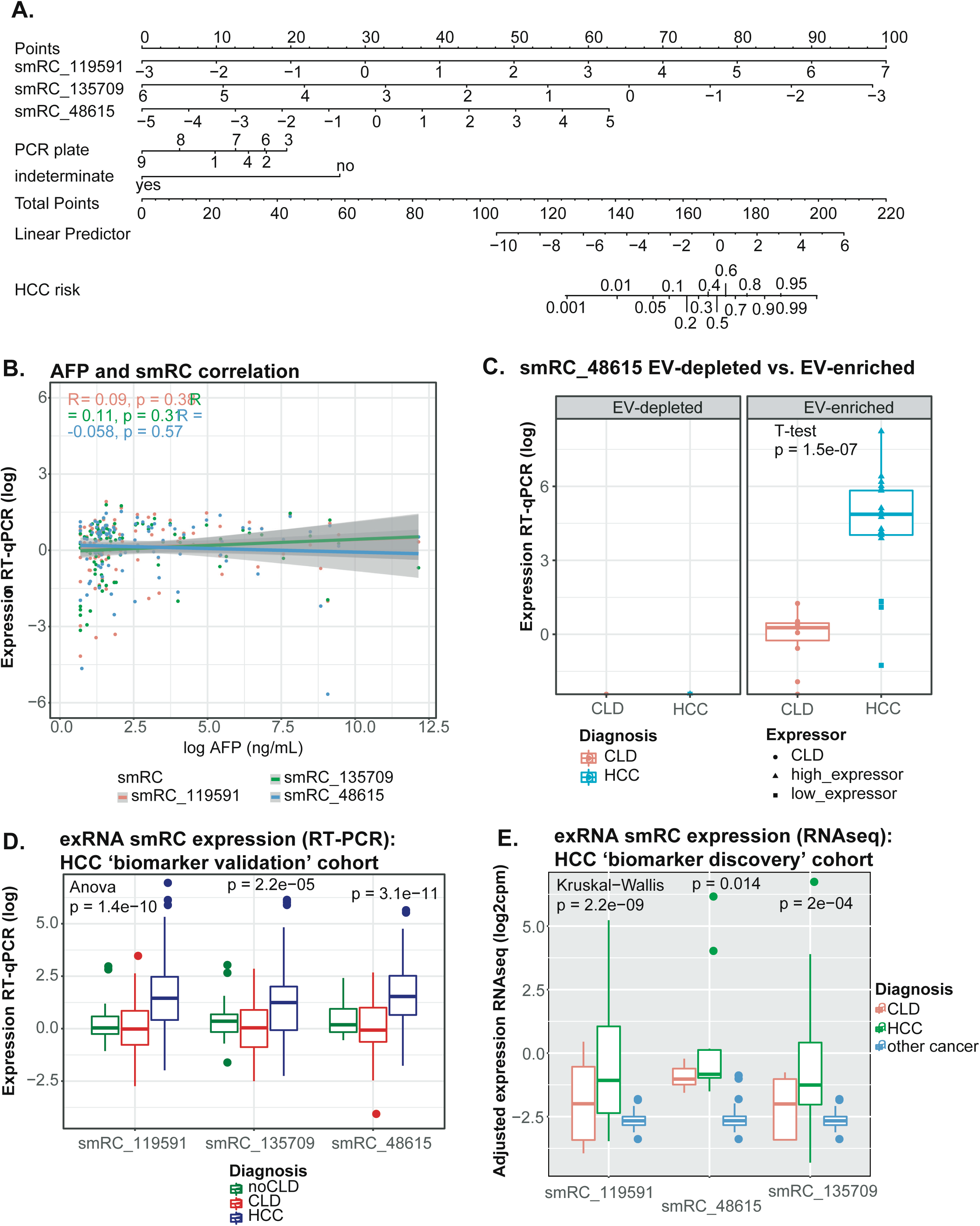
smRC in ‘HCC biomarker validation’ cohort. (**A**) Nomogram for 3-smRC signature to predict early stage HCC. (**B**) AFP and smRC correlation plot. (**C**) Expression for each smRC between HCC patients, chronic liver disease controls (CLD), and patients without chronic liver disease (noCLD). (**DC**) Expression of smRC-48615 in EV-enriched isolates and EV-depleted plasma. Displayed are samples from HCC and CLD controls. Triangles indicate HCC samples with relatively high expression, rectangles indicate samples with lower expression. (**D**) Expression for each smRC between HCC patients (n=105), chronic liver disease controls (CLD, n=85), and patients without chronic liver disease (noCLD, n=19) (RT-PCR data). (**E**) Expression for each smRC between chronic liver disease controls (CLD, n=5), HCC patients (n=10), and patients with other non-HCC malignancies (n=142) (HCC ‘biomarker discovery’ cohort, RNAseq data).

## DISCUSSION

Our study provides a conceptually novel solution to a key barrier in the field of exRNA-derived cancer biomarkers. We strongly depart from previous exRNA characterization studies, which are restricted to quantifying expression of known (i.e. annotated) transcripts^21^. Thus, we do not discard the substantial component of unannotated exRNA, or simply focus on a particular RNA biotype (e.g. miRNA)^22,23^. Instead, we provide a novel, scalable, and data-driven view of the entire small exRNA landscape unfettered by incomplete and emerging prior knowledge. This approach allowed us to identify and validate novel circulating biomarkers for the detection of curable HCC.

By *de novo* characterizing the unknown non-coding small exRNA landscape across EV isolation technologies, biofluid, and cancer type, we have defined the key properties of exRNA-associated smRCs, including their clinical application in early cancer detection. We have used a comprehensive dataset from prostate cancer patients to establish an exRNA-specific smRC feature set from which we mine their key statistical properties and develop selection criteria. These properties indicate that the tractable smRC-based quantification of novel, unannotated, small RNA expression signatures is feasible across different EV isolation techniques applied to different biofluids, potentially offering a completely novel, data-driven strategy for increasing the sensitivity of EV-driven biomarker discovery. It is important to emphasize that multiple small functional noncoding RNA can arise from transcriptional post-processing of a single larger RNA precursor gene (e.g. endogenous siRNAs of plants^24^ and animals ^25^, miRNA hairpins yielding miRNA*^26^, and piRNAs^27^), so smRCs estimate the overlooked underlying expression profile of small RNA precursor genes and thereby facilitate accurate quantification, differential expression, and motif discovery of unknown, heterogeneous, small RNA dominated exRNA payloads. In this sense, smRCs might more accurately measure the information content of exRNA. Applying our approach to a separate HCC plasma-based exRNA dataset, we derive a 3-smRC (unannotated), HCC-specific signature to discriminate patients with incipient HCC from controls at high-risk of cancer. In line with recommendations of the International Society of Extracellular Vesicles^12^, a thorough characterization of our isolates suggested a predominant enrichment for small EVs as the most likely origin of our smRC signal.

Importantly, our exRNA-derived smRC signature was developed as a method for early HCC detection in the context of cancer surveillance and not as a HCC diagnostic tool. There is a subtle but very crucial difference between these two clinical scenarios which directly determined the patient population we deliberately selected for this study, as extensively outlined in clinical guidelines^10,19^. Briefly, these guidelines explicitly underscore the urgent clinical need for new tools to detect patients with early stage HCC, as they can be cured if diagnosed at this stage. Other malignant liver tumors (e.g. cholangiocarcinoma) and associated metastases rarely occur in patients with cirrhosis and are not the target of liver cancer surveillance programs^7^. Nevertheless, we have confirmed the HCC specificity of our 3 smRC signature in a dataset of 142 patients with other malignancies. We purposely chose to test our early detection biomarker candidates in the context of the hardest possible scenario of distinguishing between chronic liver disease and very early, curable, HCC. Our signature is independently validated in more than 200 patients, where we demonstrate its ability to accurately detect patients with early stage HCC. We demonstrate that our 3-smRC signature not only outperforms the recommended surveillance tools (serum alpha-fetoprotein (AFP) combined with abdominal ultrasound)^11^, but is complementary to AFP and in combination further maximizes HCC detection rates. There are other approaches currently under evaluation for early HCC detection using other liquid biopsy analytes, mostly involving circulating DNA. Mutation^28^ and methylation^18,29^ studies have shown comparable performance to our 3-smRC signature. The main difference with our study is that most of them included HCC patients at more advanced stages^30^ as opposed to our exclusively early-stage cohort.

Despite not yet having a clear functional role in oncogenesis apart from suggestive enrichments in key RNA binding protein motifs (**Supplementary Fig. 6**), our findings strongly suggest that unannotated smRCs enable a robust, blood-based, minimally invasive, operator independent surveillance test for HCC, which is a major unmet clinical need in at-risk patients. While further validation in phase 3 biomarker studies will pave the way for its clinical implementation, this study highlights complex, heterogeneous, non-coding and unannotated small RNA payloads of EVs and their emergence as a powerful modality for biomarker discovery in cancer.

## MATERIALS AND METHODS

### Patient enrollment

For the prostate cancer dataset, de-identified data and biospecimens from human subjects consented under ongoing IRB approved protocols at the Icahn School of Medicine at Mount Sinai (GCO# 14-0318, 15-1135 and 10-1180) were collected from prostate cancer patients undergoing prostatectomy. Specifically, biospecimen included prostate cancer and adjacent prostate non-tumoral tissue from biopsy or prostatectomy, urine, and serum, where applicable. Each of these protocols involves the prospective collection of clinical data (e.g., demographics, baseline characteristics, treatments, and outcomes). Samples for the HCC ‘biomarker discovery’ and ‘biomarker validation’ cohorts were collected from consented patients enrolled in an IRB approved protocol to derive new HCC biomarkers from blood (HS-15-00540) or provided by the Tisch Cancer Institute Biorepository (HSM#10-00135) at the Icahn School of Medicine at Mount Sinai. For the phase 2 biomarker study, we included three patient populations: 1) HCC cases were limited to very early or early stage patients according to the BCLC classification^7^ (i.e., stages 0 or A). All HCC patients were treatment-naïve at the time of blood sampling, 2) Patients with liver cirrhosis or different forms of chronic liver disease (CLD) at risk for HCC as per clinical practice guidelines^10,19^, but without radiological evidence of HCC at the time of blood collection, 3) patients with benign liver nodules (e.g., hemangioma) without chronic liver disease. HCC diagnosis was made according to the criteria of the European Association for the Study of the Liver (EASL)^19^. Liver cirrhosis was diagnosed based on histology, or non-invasively through combined transient elastography, imaging or laboratory evidence of liver dysfunction and portal hypertension. Patients with concurrent malignancies were excluded. Small RNA sequencing data from patients with other (non-HCC) malignancies were downloaded from exRNA atlas (https://exrna-atlas.org/, including n=100 colon cancer, n=6 pancreatic adenocarcinoma, and n=36 prostate cancer patients, respectively).

Methods and data on **Sample collection and separation of EVs from human plasma, serum, and urine, EV characterization, RNA extraction, small library preparation and next-generation sequencing, trimming, alignement, deconvolution analysis, smRC definition and properties, smRC overlap with known biotypes** and **prostate cancer motif sequences** can be found in the supplementary material.

### Reverse transcriptase quantitative polymerase chain reaction (RT-qPCR)

We designed custom TaqMan® Small RNA Assays to target our candidate smRCs (ThermoFisher, **Supplementary Table S4+S5)** and purchased a catalog TaqMan® miRNA Assay against cel-miR-39-3p (ThermoFisher) to target the spike-in miRNA mimic which was used during the exRNA extraction. Three μl of extracted exRNA were used for reverse transcription (RT) to cDNA with the conventional TaqMan™ MicroRNA Reverse Transcription Kit (ThermoFisher) and target-specific RT primers, followed by quantitative real-time PCR according to the manufacturer’s protocol. For our 3-smRC signature, raw ct values of smRCs were corrected against ct values of the spike-in (ΔCt) and normalized to the average ΔCt of all controls (ΔΔCt). Overall, the turnaround time from blood sampling to final test results can be achieved in less than 12 hours.

### Data Analysis

Following the guidelines of the Early Detection Research Network by the National Cancer Institute^31^, we conducted a population-based case-control phase 2 biomarker study for early detection of HCC. Based on the largest meta-analysis on surveillance for HCC, sensitivity of the current gold-standard for HCC surveillance (i.e., abdominal ultrasound and AFP) is 63% for early stage tumors^11^. We powered this study to detect an increase in sensitivity from 63% to 75% and specificity from 83% to 95%. Given an alpha of 0.05 and a power (1-ß) of 80%, the number of samples needed to detect this difference based on asymptotic normal distribution theory^32^ was 101 cases (early HCC) and 71 controls (patients at high risk of HCC, CLD). For descriptive statistics, continuous variables are reported as median and categorical variables as counts and percentages. We used the Fisher’s exact test and the Student’s t-test to compare differences between categorical and continuous variables, respectively. Pearson’s or Spearman’s correlation coefficients were computed for correlation of continuous variables as indicated. Boxplot center line shows median, box limits show upper and lower quartiles, whiskers show 1.5x interquartile range, and points represent outliers. Error bars represent the 95% confidence intervals

The analysis of the phase 2 biomarker study to test the performance of the 3-smRC signature for early detection of HCC was limited to early stage HCC (n=105) and controls at risk for HCC (CLD, n=85) to represent the optimal population of interest.^17,19^ We used penalized maximum likelihood techniques, bootstrap and cross-validation to estimate and control for model optimism, RT-qPCR batch plate effects, and over-fitting, and also rigorously computed the positive and negative predictive power estimates of our 3-smRC early detection signature. We computed a number of indices of model performance, discrimination measures, and calibration measures under bootstrap resampling (n = 1000), as summarized in **Fig. 4D**, in order to demonstrate model performance and estimate generalization error by averaging performance across bootstrap resampling. In the first row, the key measure of discrimination Somers’ Dxy is the rank correlation between the observed and predicted response values, which in the case of logistic regression for a binary response reduces to simply Dxy = 2(c - ½), where c is Harrel’s c-statistic and equal to the AUC of the ROC for the early HCC vs. CLD prediction. In the case of the smRC model we immediately deduce that the bootstrap adjusted AUC is ½ + 3/8 = 7/8 = 0.875. Modest adjusted modified R2 ~ 0.52 is observed, combined with bootstrap-adjusted slope and intercept indicating modest and acceptably low over-fitting. Relatively bootstrap-adjusted low Emax (the maximum error in predicted probabilities), modest Brier score (B), very low unreliability index (U), high discrimination (D), high quality (Q = D - U), also indicate a reasonably robust model. Also, the bootstrap adjusted total Gini’s mean difference for based on the smRC model is a healthy 2.44, which robustly represents typical log-odds differences between early HCC and CLD patients predicted by the model. Converting this early HCC log-odds estimate to an early HCC probability prediction, we see that the typical predicted probability gap between early HCC and CLD patients is 38%. Finally, we compute the partial mean gini-scores of the smRC model predictors and find that the smRCs themselves have by far the largest termwise log-odds compared to any technical variance covariates (e.g., batch). We note in passing that repeated cross-validation gave similar results for Dxy and adjusted Slope (**Fig. 4D**, extended in **Supplementary Table S5**).

We next repeated the penalized maximum likelihood estimation procedure using a model with both smRCs and AFP readings included, given that a log likelihood ratio test for an AFP term was highly significant (p < 1e-8). Computing the same indices of model performance across bootstrap resampling (n = 1000), we found dramatically better performance as shown in **Fig. 4D**, with bootstrap adjusted AUC ~ 0.93, lower overall error and evidence for overfitting, a much smaller Brier score of 0.11, and a dramatic increase in the Gini indices such that a typical early-HCC - CLD predicted probability difference was 43% (**Fig. 4D**, extended in **Supplementary Table S5**).

Finally, even though balanced accuracy is not a proper scoring rule, we estimated the maximized balanced accuracy landscape by subjecting the smRC logistic regression model for HCC risk to a cross-validation repeated 1000 times (i.e., a random 85% training, 15% testing split repeated 1,000 times) and computing maximizing sensitivity and specificity on the test ROCs. We found strong evidence to suggest that sensitivity ~ 86% and specificity ~ 91% for smRC-only models, while for smRC + AFP models we found sensitivity ~ 85% and specificity ~ 94%.

All statistical analyses were conducted on Rstudio (R version 3.5.0).

## SUPPLEMENTARY MATERIALS

**Supplementary Fig. S1.**
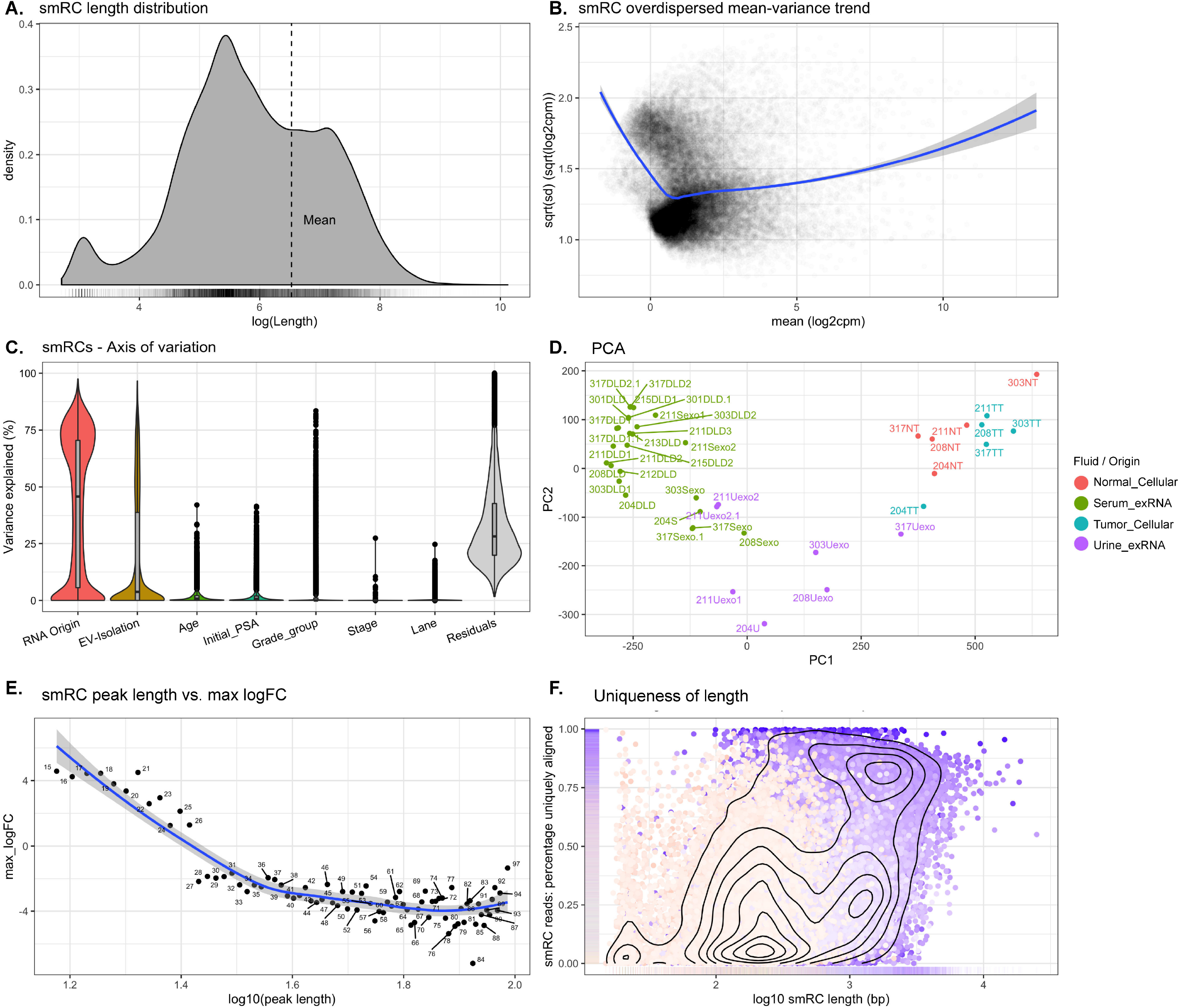
smRC properties of prostate cancer ‘smRC characterization cohort’. (**A**) Density plot of smRC length. (**B**) Mean-variance profile across all samples. (**C**) smRC axis of variation. The relative contribution of each axis of expression variation is displayed across the training prostate cancer dataset in order of magnitude. RNA origin (i.e., EV-derived or cellular) contributed the most to the observed variance. (**D**) Principal component analysis (PCA). (**E**) Maximum value logFC among all significant smRCs as a function of the length of the smRC peak consensus sequence. (**F**) Mapping uniqueness of smRC as a function of the length.

**Supplementary Fig. S2.**
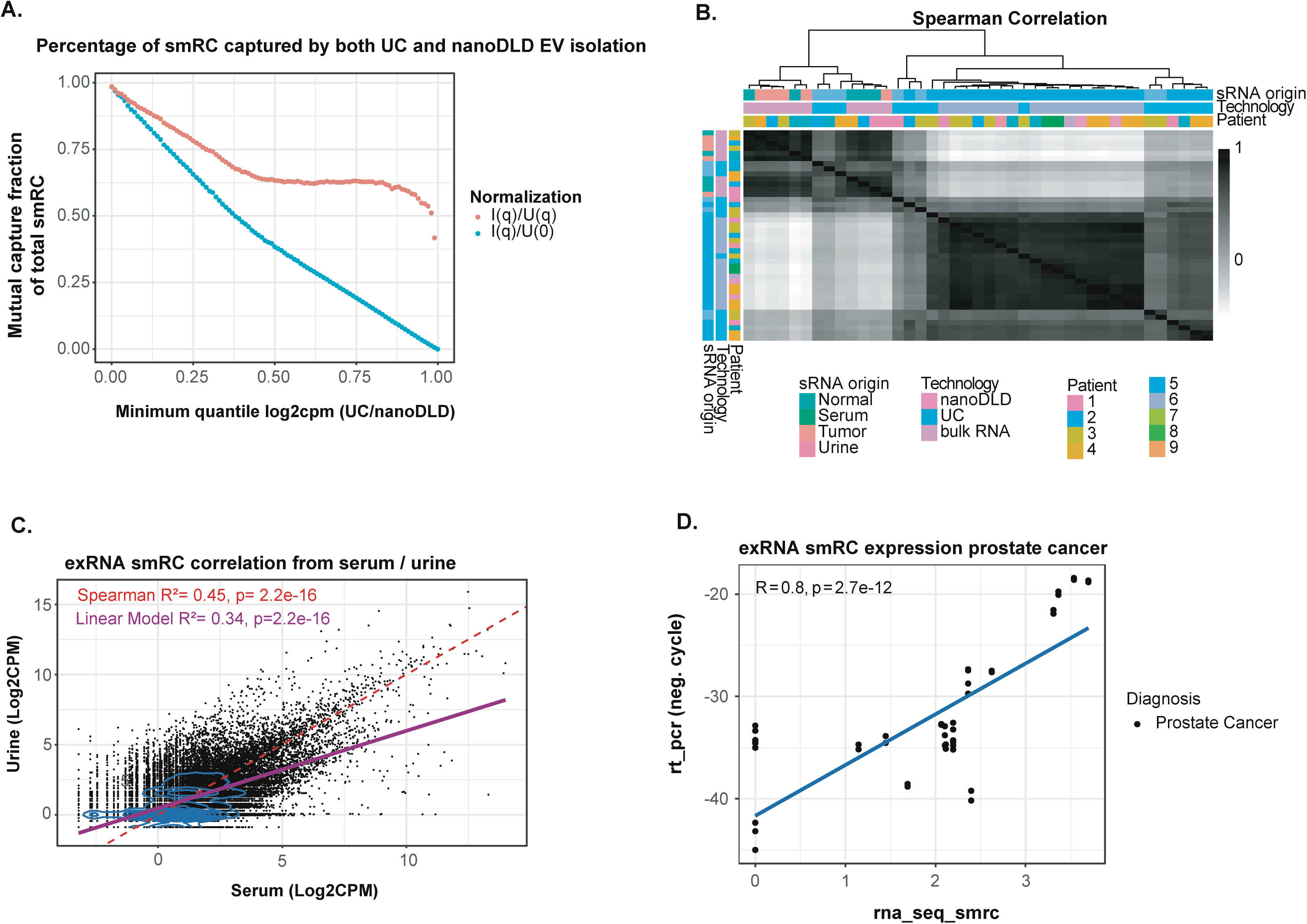
smRC correlation properties across different biofluids and technologies. (**A**) Percentage of smRC captured by both UC and nanoDLD EV isolation. (**B**) Correlation plot across prostate cancer samples. (**C**) Correlation plot for EV-derived smRC expression across different biofluids (i.e., serum *versus* urine) using UC. (**D**) Correlation of single smRC expression between RNAseq and RT-PCR in the prostate cancer cohort.

**Supplementary Fig. S3.**
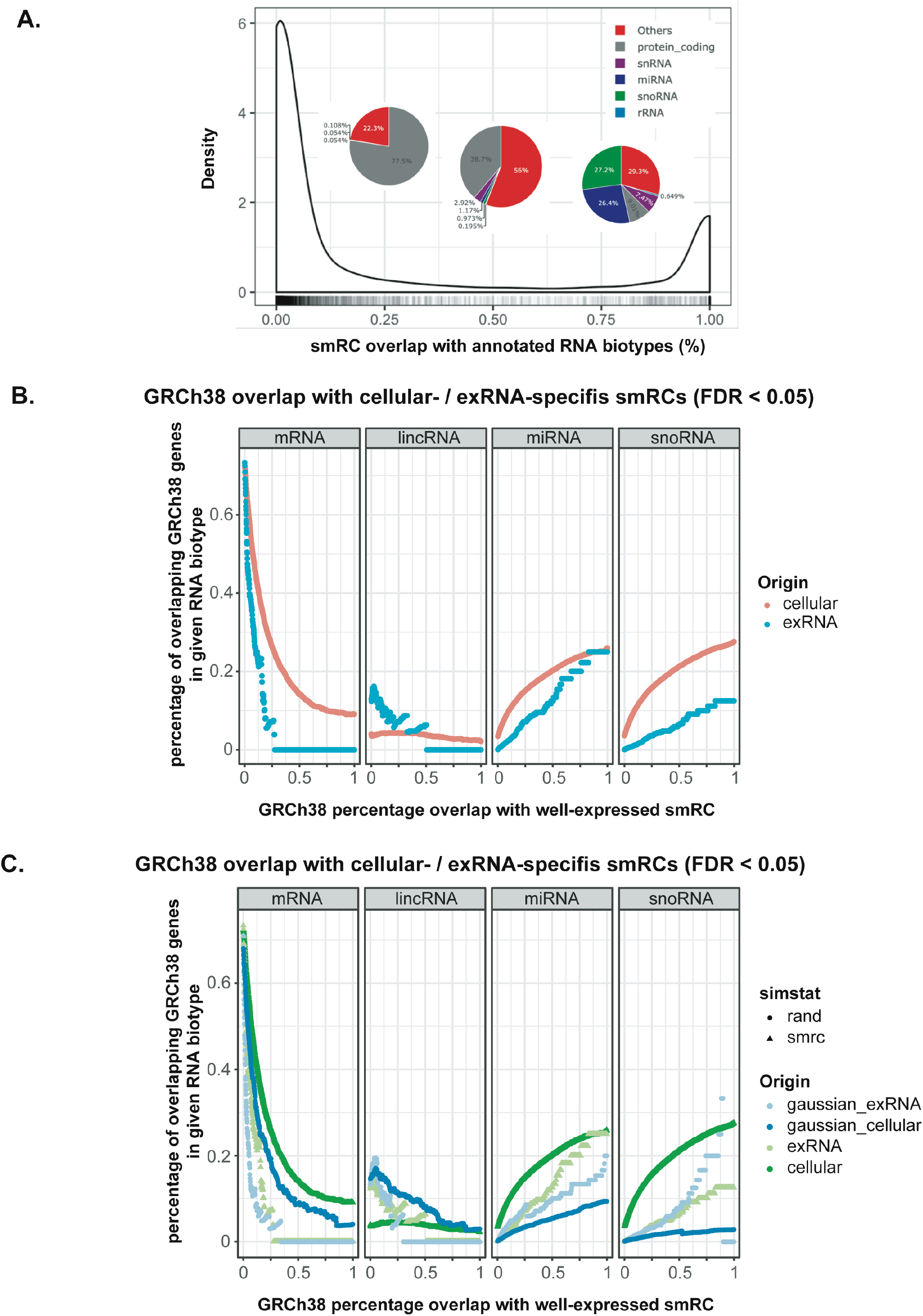
Level of smRC overlap with annotated hg38 biotypes. (**A**) Distribution of percentage overlap of smRCs onto all known hg38 RNA biotypes. Low overlap (≪1) indicates smRC does not contain whole RNA bioptype, high or total overlap (~1) indicates RNA biotype contained within smRC. (**B**) Plot of given RNA biotype abundance percentage (among all RNA biotypes in hg38 annotation) versus smRC overlap percentage as above. Abundance percentage quantifies the frequency of a given RNA biotype among all others. (**C**) Same as (B), only with curves derived from a *random* genomic distribution matching number and size of smRCs.

**Supplementary Fig. S4.**
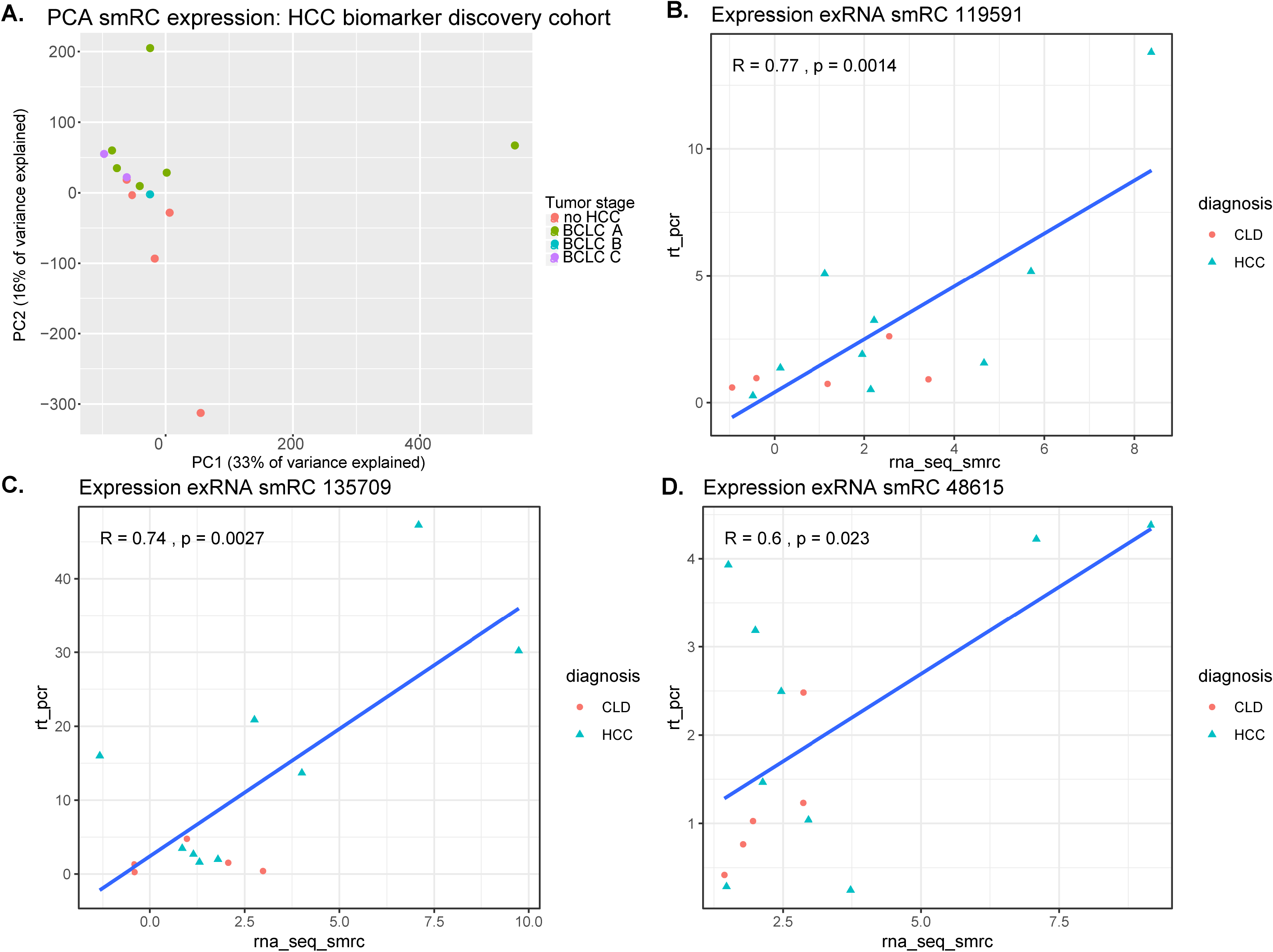
smRC in ‘HCC biomarker discovery’ cohort. (**A**) Principal component analysis (PCA) for HCC biomarker discovery cohort. (**B-D**) Correlation of 3-smRC-signature expression between RNAseq and RT-PCR in the HCC discovery cohort.

**Supplementary Fig. S5.**
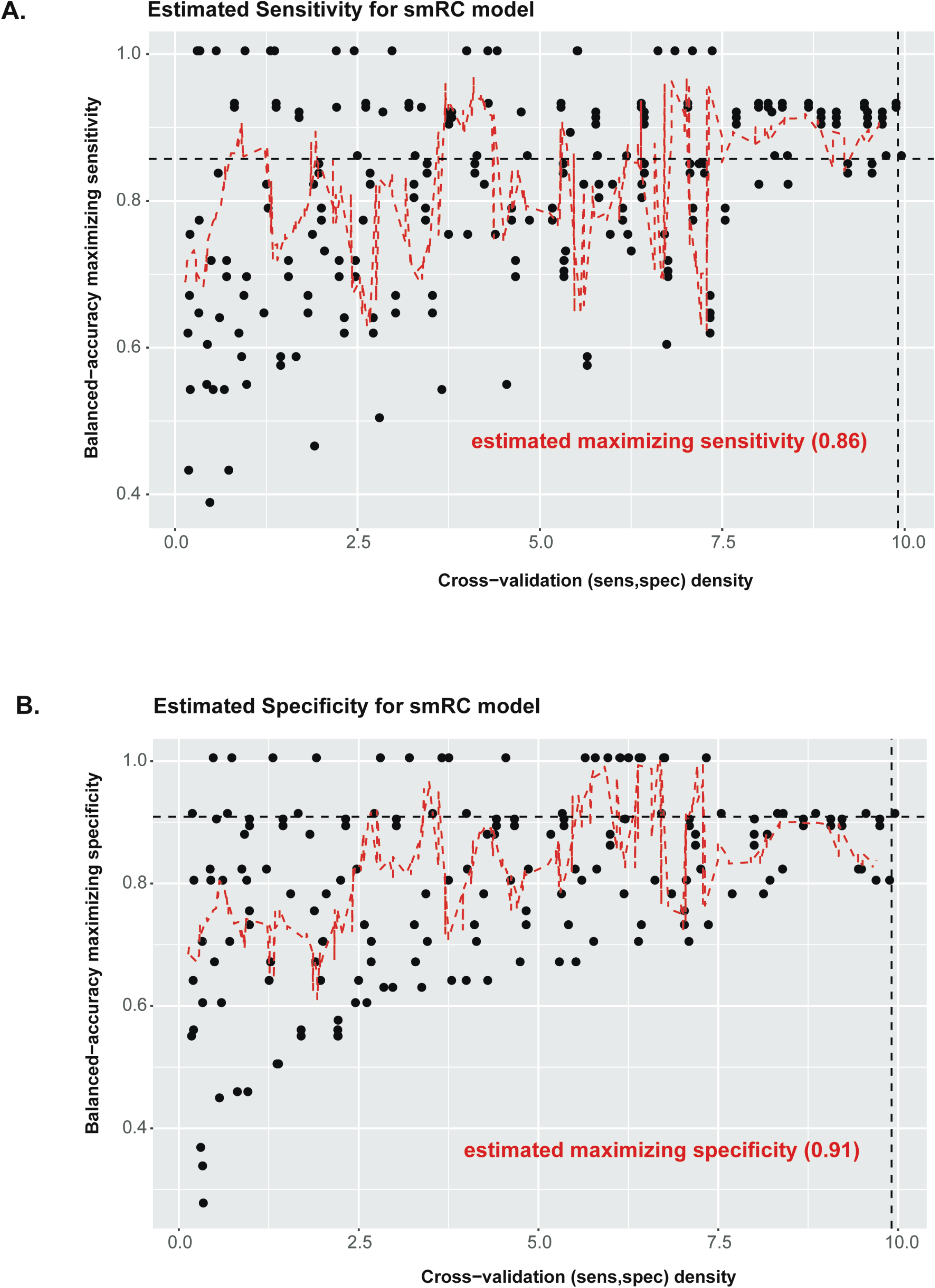
Sensitivity and specificity of smRC model. `Balanced accuracy−maximizing sensitivity` (**A**) and specificity (**B**), respectively, *versus* kernel density estimation of all [sens, spec] simulation pairs (with n = 30 moving average) for the smRC model to discriminate early stage HCC from controls at high risk.

**Supplementary Fig. S6.**
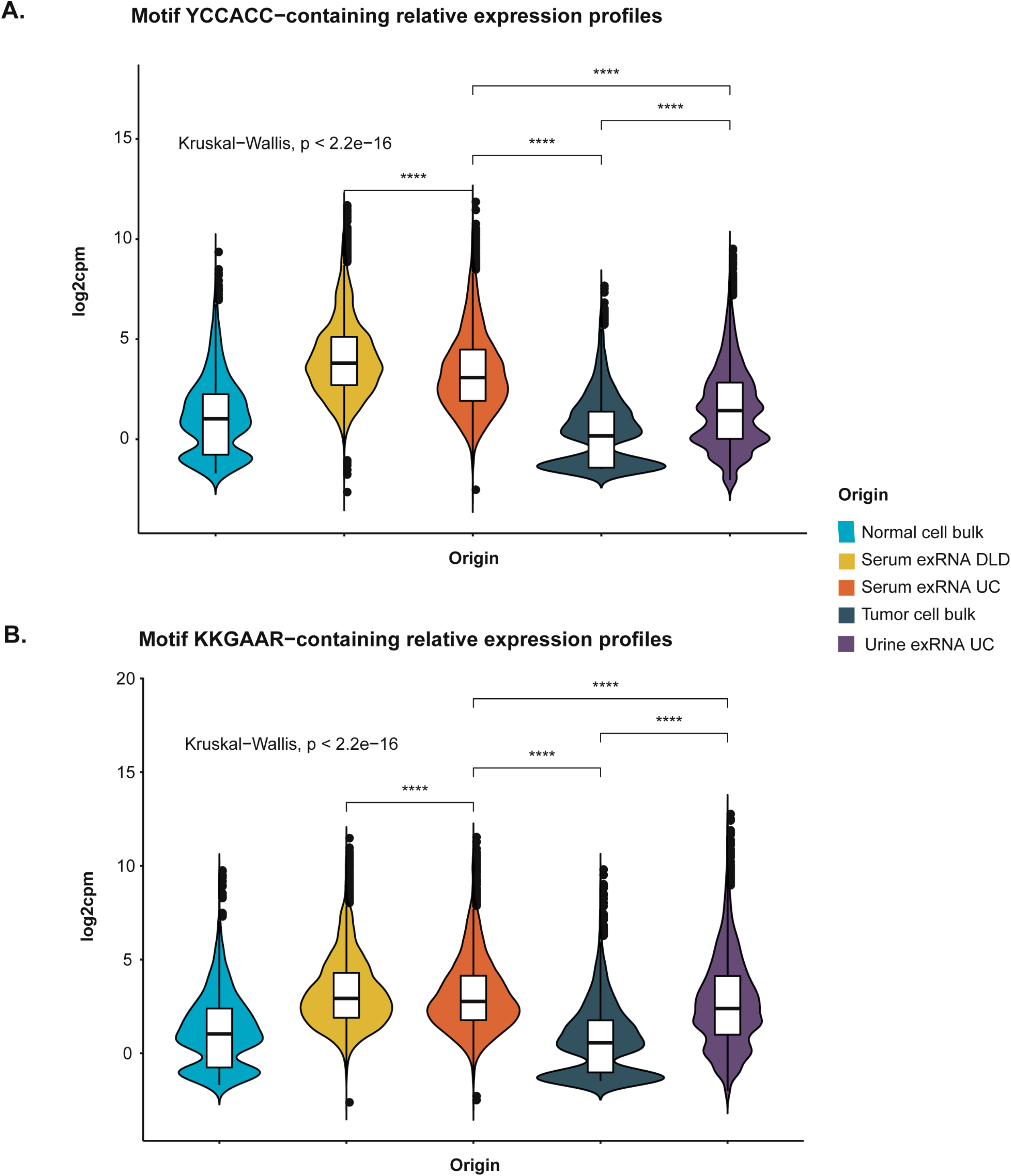
Motif containing smRC expression in prostate cancer ‘smRC characterization’ cohort across RNA origin. Expression profile of exRNA smRCs enriched in either of the motifs (**A**, YCCACC, **B**, KKGAAR) reveals over-expression in exRNA compared to cellular smRCs (true by definition) with a bimodal downregulated expression of motif-enriched cellular smRCs.

**Supplementary Fig. S7. Complete image of Western Blotting analysis targeting TSG101.** The section included in the manuscript is highlighted in red.

**Supplementary Table S1.**

| **RNA biotype** | **Biofluid** | **statistic** | **P-value** | **Alternative hypothesis** |
| --- | --- | --- | --- | --- |
| **mRNA** | cell | 0.527 | 0 | CDF x above CDF y |
| **mRNA** | exRNA | 0.106 | 1.32e-05 | CDF x above CDF y |
| **lincRNA** | cell | 0.00 | 1 | CDF x above CDF y |
| **lincRNA** | exRNA | 0.250 | 0 | CDF x above CDF y |
| **miRNA** | cell | 0.913 | 0 | CDF x above CDF y |
| **miRNA** | exRNA | 0.393 | 0 | CDF x above CDF y |
| **snoRNA** | cell | 1.000 | 0 | CDF x above CDF y |
| **snoRNA** | exRNA | 0.158 | 0 | CDF x above CDF y |
| **mRNA** | cell | 0.008 | 1 | CDF x below CDF y |
| **mRNA** | exRNA | 0.068 | .0098135 | CDF x below CDF y |
| **lincRNA** | cell | 0.689 | 0 | CDF x below CDF y |
| **lincRNA** | exRNA | 0.058 | 0.03455966 | CDF x below CDF y |
| **miRNA** | cell | 0 | 1 | CDF x below CDF y |
| **miRNA** | exRNA | 0.068 | 0.0098135 | CDF x below CDF y |
| **snoRNA** | cell | 0 | 1 | CDF x below CDF y |
| **snoRNA** | exRNA | 0.245 | 0 | CDF x below CDF y |

**Supplementary Table S2.** Clinical characteristics of the discovery cohort for HCC patients and controls.

|  | **Early Stage HCC**<br>**(n=10)** | **CLD, risk for HCC**<br>**(n=5)** | **P-Value** |
| --- | --- | --- | --- |
| **Age (Years)** | 67 | 63 | 0.94 |
| **Sex (Male)** | 7 (70%) | 3 (60%) | 1 |
| **Cirrhosis (Yes)** | 8 (80%) | 4 (80%) | 1 |
| **Etiology** |  |  |  |
| **HCV** | 4 (40%) | 2 (40%) | 1 |
| **HBV** | 3 (30%) | 1 (20%) | 1 |
| **NASH** | 3 (30%) | 2 (20%) | 1 |
| **Tumor stage (BCLC)** |  |  |  |
| **Early Stage (Stage A)** | 6 (60%) | n.a. | n.a. |
| **Intermediate Stage (BCLC B)** | 2 (20%) | n.a. | n.a. |
| **Advanced Stage (BCLC C)** | 2 (20%) | n.a. | n.a. |
| **Largest nodule (cm)** | 3.5 | n.a. | n.a. |
| **AFP (ng/mL\*)** | 20.6 | 4.6 | 0.46 |

Continuous variables are displayed as median. \*Upper limit of normal 9ng/mL. AFP, alpha fetoprotein, BCLC, Barcelona Clinic for Liver Cancer, HBV/HCV, chronic hepatitis B/C, NASH, non-alcoholic steatohepatitis

**Supplementary Table S3.**

| **Supplementary Table S3** |  |  |
| --- | --- | --- |
| **Read mapping status** | **PrCA smRC**<br>**characterization cohort**<br>**(reads)** | **HCC smRC biomarker**<br>**discovery cohort (reads)** |
| **Total (post cutadapt)** | 494 828 430 | 409 592 240 |
| **Unmapped** | 213 988 996 (43.2%) | 138 477 962 (33.8%) |
| **Unmapped because m > 50** | 10 370 278 (2.1%) | 11 205 827 (2.7%) |
| **Unmapped because guidance failed** | 38 714 298 (7.8%) | 7 066 592 (1.7%) |
| **Uniquely mapped (U)** | 86 369 038 (17.5%) | 203 895 810 (49.8%) |
| **Multiply mapped (m < 3, R)** | 2 136 572 (0.4%) | 2 072 756 (0.4%) |
| **Multiply mapped with u-rescue (P)** | 143 249 248 (28.9%) | 46 873 293 (11.4%) |
| **Primary alignments (U + R + P)** | **229 618 286 (46.4%)** | **250 769 103 (61.2%)** |

Supplementary Table S4. RT-qPCR assay sequences of 3-smRC signature and genomic location.

**Supplementary Table S5.**
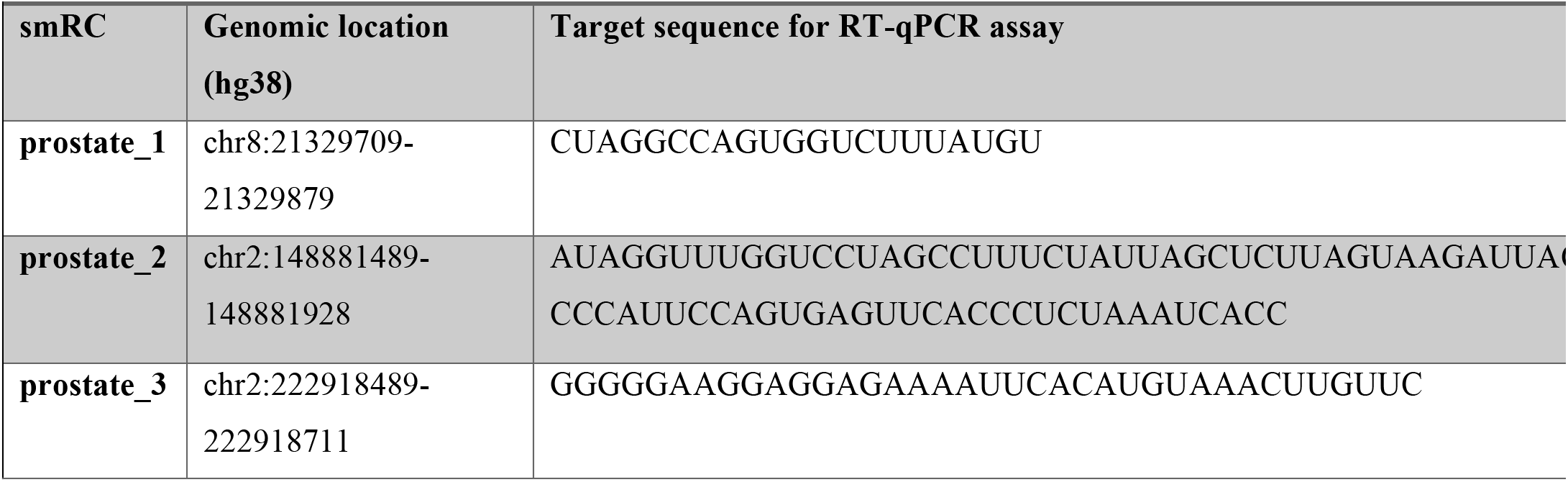
RT-qPCR assay sequences for orthogonal smRC validation in prostate cancer dataset

**Supplementary Table S6.**
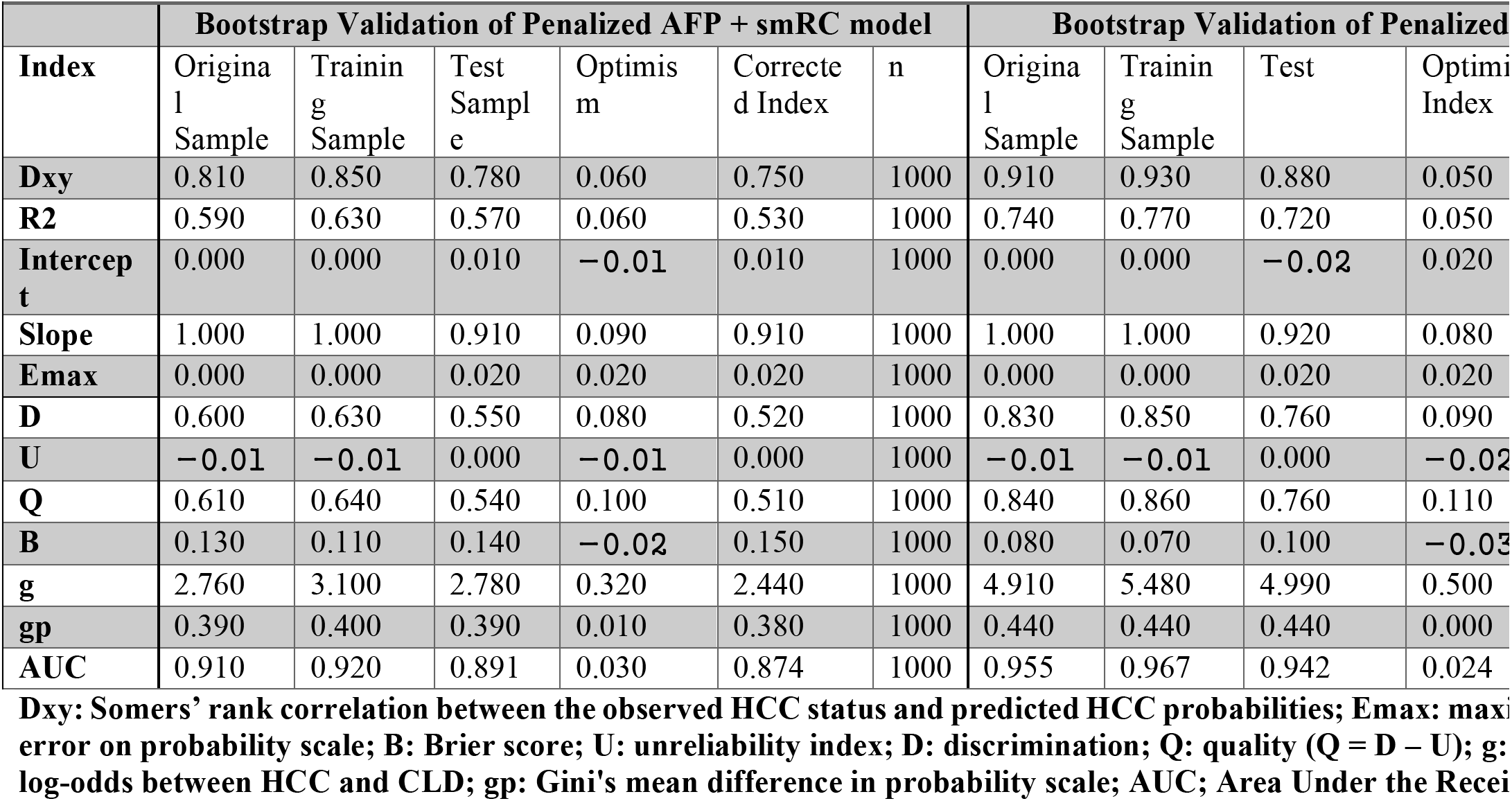

## ACKNOWLEDGEMENTS

The authors thank the office of Scientific Computing and the Genomics Core Facility at the Icahn School of Medicine at Mount Sinai (ISMMS) for providing computational resources and staff expertise, as well as the ISMMS Tissue Biorepository for providing some of the samples. The authors further thank Dr. Veronica Sanchez-Gonzalez (NanoView Biosciences, Boston, MA) for helping with the Exoview™ analysis.

## SUPPLEMENTARY METHODS

### Sample collection and separation of EVs from human plasma, serum, and urine

For the prostate cancer dataset, human serum was collected using BD Vacutainer blood collection tubes (i.e., serum separation tubes). First, whole blood was centrifuged at 2,000g for 30 minutes at 4°C followed by another centrifugation of the serum at 12,000g for 45 minutes at 4°C to remove larger EVs (e.g. microvesicles and apoptotic bodies). The supernatant was carefully transferred to ultracentrifugation tubes (Beckman coulter, thick wall polypropylene tube, Cat. #355642) and ultracentrifuged for two rounds at 110,000g for 2 hours at 4°C. The pellet was finally resuspend in 1 mL PBS and stored at −80C for further analysis. EVs from human urine were collected with the above mentioned protocol.

For the HCC ‘biomarker discovery’ and ‘biomarker validation’ dataset, peripheral venous blood was collected in EDTA containing vacutainer (BD Vacutainer), stored on ice, and processed within 4 hours of collection. On the day of collection, we performed two centrifugation steps to separate plasma from other blood components and minimize cellular debris from our final isolate. First, whole blood was centrifuged at 1,600g for 10 minutes at 4°C followed by another centrifugation of the plasma at 16,000g for 10 minutes at 4°C to remove larger EVs (e.g. microvesicles and apoptotic bodies). The supernatant was then stored at −80°C until the ultracentrifugation was performed. For this, samples were thawed on ice and 0.5 - 1 mL of plasma was diluted in ~25 mL PBS and centrifuged at 120,000g for 2 hours at 4°C with a Type 50.2 Ti Fixed-Angle Titanium rotor (Beckman Coulter, k-factor = 69). Isolates were directly used for RNA extraction (see below) or resuspended in PBS and stored at −20°C until further analysis.

### EV characterization

EV characterization procedures followed the recommendations by the International Society for Extracellular Vesicles (ISEV)^12^. After differential ultracentrifugation, the PBS-resuspended isolate was evaluated with transmission electron microscopy (TEM) in a Hitachi 7000 transmission electron microscope operating at 80 kV. Briefly, equal volumes of the isolate and 3% Glutaraldehyde were mixed and kept at room temperature for 1 hour. Two μl of osmium tetroxide was added to the mixture and incubated at room temperature for 1 hour. The solution was then transferred to formvar coated TEM grids and observed under the electron microscope. To estimate the size and concentration of the isolate, we conducted nanoparticle tracking analysis (NTA) on a NanoSight NS300 (Malvern Instruments Ltd, Malvern, UK) and analyzed the samples with the NTA 3.2 software (Malvern). For this, PBS-resuspended isolates were diluted 1:50 in PBS.

For immuno-labeling of the isolate, we performed Western Blotting for the intracellular marker TSG101 and Exoview™ analysis for colocalization of tetraspanins CD9, CD63, and CD81. For Western Blotting, we quantified protein concentration (Bradford assay, Biorad) and 20 μg of protein were separated by sodium dodecyl sulfate-polyacrylamide electrophoresis under reducing conditions and transferred to PVDF membranes (Life Technologies). Unspecific binding sites were blocked with 5% nonfat dry milk and membranes were incubated with mouse monoclonal TSG101 antibody (ab83, Abcam) at 4°C overnight followed by goat anti-mouse secondary antibody (A0447, Agilent Technologies) for 1 hour at room temperature. Chemiluminescence was detected using the ECL™ Prime Western Blotting System (RPN2232, GE Healthcare). The uncropped Western Blot image for TSG101 is displayed in **Supplementary Fig. S7**. Exoview™ experiments were carried out on an ExoView™ R100 imaging platform (NanoView Bioscience). With the Exoview™ Tetraspanin kit, 35 μl of PBS-resuspended isolate was incubated overnight on a microarray chip which has been functionalized with antibodies against CD9, CD63, CD81, plus IgG negative control to detect EVs expressing these surface markers. After washing off unbound particles, chips were stained with fluorescence-conjugated antibodies against CD9 (Alexa 647) or CD81 (Alexa 555) to identify subpopulations based on maker profiles. Analysis was done with the NanoViewer 2.4.5 (NanoView Bioscience).

### RNA extraction, small library preparation and next-generation sequencing

For the prostate cancer dataset, total RNA was extracted from the serum/urine bump fraction (nanoDLD, serum only), UC isolates, or bulk tissue using the Total Exosome and Protein Isolation Kit (Invitrogen 4478545) by following the protocol. For the HCC biomarker discovery and biomarker validation datasets, RNA was extracted from the UC isolate on the same day of ultracentrifugation using the miRNeasy Plasma/Serum kit (Qiagen) according to the manufacturer’s recommendations including the spike-in *C. elegans* miR-39 miRNA mimic and stored at −80°C until further use. RNA quantitation and quality was assessed on a 2100 Bioanalyzer Instrument (Agilent) with the RNA 6000 Pico Kit (Agilent). Indexed Illumina Small RNA libraries were prepared with the SMARTer® smRNA-Seq Kit (Clontech Laboratories, Inc.) and sequenced on an Illumina HiSeq 4000 (prostate cancer dataset) or HiSeq2500 (liver cancer dataset) platform.

### Trimming

The SMARTer™ smRNA-Seq kit yields reads are flanked on the 5’ end by a leading triad of three bases from SMARTer™ template switching activity, and on the 3’ end by the Illumina adapter and extra bases from the oligo dT (which are exactly 15 bp in length). We used Cutadapt^33^ to remove the first 3 nucleotides of all reads, specify the homopolymer adapter sequence AAAAAAAAAA to remove along with any sequence 3’ of it, and finally discard all reads that are smaller than 15 bp long after these filters are applied. The exact command used, as recommended by the (strand-sensitive) SMARTer™ smRNA-Seq kit, is

~~~
cutadapt -m 15 -u 3 -a AAAAAAAAAA input.fastq > output.fastq
~~~

Therefore our set of initial small RNAs are at least 15 bp long, and are trimmed from positions 1-3 and also from the oligo dT 3’ through to the adapter. We note in passing that although template switching at low frequencies can add more than 3 nucleotides to the 5’ end, we did not trim any further on the 5’ end.

### Deconvolution analysis

EV carrier deconvolution analysis was performed as a post-processing step to the standard exceRpt pipeline^40^, which was applied to the entire HCC smRC discovery dataset (n=15). The output of exceRpt is collated (using mergePipelineRuns.R from https://github.com/rkitchen/exceRpt) to form summary data of count matrices for key annotated, noncoding RNA biotypes (piRNA, circRNA, miRNA, tRNA counts), aggregated QC data, adapter sequence data, and diagnostic plots. At this point we applied their deconvolution algorithm on the summarized data. Briefly, this consists of two key stages: In the first stage, constituent cargo profiles are estimated using a modified version of a methylation deconvolution technique in Onuchic et al.^41^. Next, deconvolution is performed using the Read Counts or RPM sample profiles from the exRNA Atlas and the per-sample proportion enrichments of each profile are estimated.

### smRC overlap with known RNA biotypes

We next investigated if well-expressed exRNA and cellular smRCs preferentially capture (enclose) any key known RNA biotypes, as we would expect with both exRNA and cellular smRCs for miRNA for example, and to what extent they do so across all key biotypes. Indeed, for a specific RNA biotype we first computed the smRC capture percentage (i.e., whether or not the smRC completely or only partially enclosed the RNA biotype). Then, for a particular smRC capture percentage, we asked how frequent a particular RNA biotype was among all biotypes. **Supplementary Fig. S3A** shows the relative breakdown of RNA biotypes at several extremal points of the smRC capture percentage (1%, 70%, 100%), where plainly miRNA, snoRNA, snRNA, and other small RNA are preferentially *completely* captured (i.e., they are the dominant RNA biotypes with capture overlap ~ 1) by smRCs compared to mRNA, which are dominantly *grazed* (i.e., protein coding biotype is dominant for capture overlap ≪ 1). In other words, as expected, when a smRC completely or mostly encloses a known RNA biotype, it is mostly likely a small RNA and very unlikely a protein-coding RNA. Indeed, plotting the RNA biotype frequency across all exRNA and cellular smRC capture overlap percentages separately for mRNA, lincRNA, miRNA, and snoRNA, yields **Supplementary Fig. S3B**. We find that exRNA smRCs dominantly *partially capture (graze)* mRNA at most to about 25% of the mRNA transcript, and never capture more, while cellular smRCs tend to overlap more protein-coding mRNA and can actually completely enclose mRNA. Similarly, using **Supplementary Fig. S3B**, one can conclude miRNA are preferentially completely enclosed by both exRNAEV and cellular smRCs at the same rate, at most 50% of a lncRNA is captured by an exRNA smRC, and snoRNAs are preferentially completely enclosed by cellular smRCs compared to exRNA smRCs. Taken together, when smRCs do enclose known RNA biotypes they can either do so predominantly partially (as with mRNA and lncRNA) or predominantly completely (as with miRNA, snoRNA, and other small RNA), with key differences in the statistics observed between exRNA and cellular smRCs. Finally, one can ask if these overlap properties are principally driven by the number and relative size distributions of exRNA and cellular smRCs (as opposed to a genuine property of small RNA accumulation in exRNA and cells). Randomly generating genomic regions with the same number of regions, and exRNA and cellular smRC size distributions (masking for repeat regions and centromeres), we repeat the above overlap computations and use a Kolmogorov-Smirnov test to assess if the underlying distributions of overlaps and capture percentages are the same within sampling noise. It turns out that all pairwise (x = smRC, y = random) Kolmogorov-Smirnov tests with two sided alternative hypotheses are highly significant, especially for lncRNAs, indicating that the exRNA and RNA biotype specific overlap patterns are not solely attributable to the size distributions (or number) of smRCs. As **Supplementary Table S1** illustrate for the two separate one-sided KS tests, interesting trends emerge: for mRNA, both exRNA and cellular smRCs tend to overlap more exons than expected by random simulation; for lncRNA, exRNA smRCs overlap than expected more while cellular smRCs overlap much less; for miRNA, both exRNA and cellular smRCs overlap far more than expected by chance; for snoRNA, cellular smRCs overlap far more than expected while exRNA smRCs have slightly more evidence for relative depletion.

In summary, exRNA smRCs overlap known RNA biotypes in a non-random fashion, and when they completely or almost completely enclose a biotype it is overwhelmingly likely to be a known *small* RNA biotype, as opposed to similar but distinct trends for cellular smRCs. To aid in interpretation and comparison, **Supplementary Fig. S3C** also includes the simulated fractional overlap curves. However, as **Supplementary Fig. S3A** demonstrates *a significant fraction of exRNA smRCs are well-expressed from unannotated genomic regions*.

### Prostate smRC consensus sequence motifs

In the absence of functional data, we speculate that like other small RNA, exRNA small RNA payloads are in complex with RNA binding proteins (RBPs), or may bear vestigial evidence of exRNA related packing by RBPs. Using MEME^44^, we investigated if the exRNA smRC peak consensus sequences had any evidence of being enriched in ungapped motif sequences that in turn had homology to known RBP motifs. Parsimoniously, we assumed that each peak sequence contains at most **one** occurrence of a motif, but likely none. We also assumed that if nucleotide frequency biases exist there would be only single-nucleotide biases (as opposed to dimer biases such as GC content, or even higher order biases), and only searched for motifs between 3 and 6 nucleotides long, rejecting all those that had a sufficiently high E-value (probability of being found randomly). This amounts to running an instance of a zeroth order Hidden Markov Model in the zoops (zero or one per sequence) setting of MEME on a fasta file of exRNA-specific smRCs, which we took to be those with positive logFC in the null H0 and FDR < 0.001:

~~~
meme exRNA_smRC_peaks.fasta -brief 100000 -rna -oc exRNA_smRC_output -nostatus -evt .001 -mod zoops -nmotifs 10 -minw 3 maxw 6 -objfun classic -markov_order 0 1> stdout 2> stderr
~~~

The final results of the MEME computation are summarized here. Briefly, two 6 nucleotide motifs were found significantly over-enriched in two distinct groups of exRNA smRC peak consensus sequences, each representing approximately 11% of the total number of exRNA smRC peak consensus sequences interrogated. The motifs YCCACC (617 smRC peaks, RBP binding prediction: PCBP1, G3BP1, HNPRL, YBX1, ELAV1, E-value ~ 1e-46) and KKGAAR (626 smRC peaks, RBP binding prediction: ESRP2, HNRPRF, HNRPH1-3, SRSF1, E-value ~ 1e-8) were submitted for RBP motif homology assessment using ATtRACT^45^ (https://attract.cnic.es/searchmotif) and RBPDB^46^ (http://rbpdb.ccbr.utoronto.ca/).

Examining the expression profile of exRNA smRCs enriched in either of these motifs across the prostate cancer ‘smRC characterization’ cohort reveals over-expression in exRNA compared to cellular smRCs (true by definition), for example in **Supplementary Fig. S6**, but interesting sub-patterns emerge. These include a bimodal downregulated expression of motif-enriched cellular smRCs, suggesting an enriched subset that might imply a role in exRNA packing within cells, and an overall upregulation in nanoDLD isolation compared to UC within serum (and also overall compared to urine UC).

